# A live-cell imaging system for visualizing the transport of Marburg virus nucleocapsid-like structures

**DOI:** 10.1101/658419

**Authors:** Yuki Takamatsu, Takeshi Noda, Stephan Becker

## Abstract

Live-cell imaging is a powerful tool for visualization of the spatio-temporal dynamics of living organisms. Although this technique is utilized to visualize nucleocapsid transport in Marburg virus (MARV)- or Ebola virus-infected cells, the experiments require biosafety level-4 (BSL-4) laboratories, which are restricted to trained and authorized individuals. To overcome this limitation, we developed a live-cell imaging system to visualize MARV nucleocapsid-like structures using fluorescence-conjugated viral proteins, which can be conducted outside BSL-4 laboratories. Our experiments revealed that nucleocapsid-like structures have similar transport characteristics to nucleocapsids observed in MARV-infected cells. This system provides a safe platform to evaluate antiviral drugs that inhibit MARV nucleocapsid transport.

## Introduction

Marburg virus (MARV), together with the Ebola virus (EBOV), belongs to the family *Filoviridae*, and has a roughly 19 kb non-segmented, single-stranded, negative-sense RNA genome. It causes severe hemorrhagic fever with high fatality rates. MARV epidemics have occasionally been reported in Central Africa, with the largest one, having a 90% fatality rate, being reported in Angola between 2004 and 2005 (CDC, 2014). Currently, there are no approved vaccines or antiviral therapeutics available to prevent or treat MARV infection. Therefore, understanding the interplay between viral and host proteins during MARV replication is necessary to establish countermeasures for the diseases. For example, revealing the mechanisms for the assembly and transport of nucleocapsids, which comprise viral genomic RNA, nucleoprotein (NP), viral proteins (VP24, VP30, VP35), and polymerase L, and are responsible for the transcription and replication of the viral genome, might contribute to the development of new therapeutic options.

The main nucleocapsid protein of MARV is NP, which is responsible for the encapsidation of single-stranded viral genomic RNA (Bharat *et al*., 2012, Muhlberger *et al*., 1999). In addition to NP, MARV nucleocapsids also contain the minor matrix protein VP24 and the polymerase cofactor VP35, both of which are essential structural elements that directly interact with NP to build a helical nucleocapsid approximately 1000 nm in length and 50 nm in diameter (Huang *et al*., 2002, Watanabe *et al*., 2006, Bharat et al., 2012). Furthermore, the viral polymerase L and the transcription factor VP30 are also associated with the nucleocapsid (Hartlieb *et al*., 2003, Biedenkopf *et al*., 2013). The core complex of the nucleocapsid, formed by the NP, VP35, and VP24 proteins together with the viral RNA, is defined as a nucleocapsid-like structure (NCLS, Fig. 1a). In MARV- and EBOV-infected cells, immunofluorescence microscopy, as well as live-cell imaging, have been used to visualize and analyze nucleocapsids by fluorescently labeling the nucleocapsid proteins (Schudt *et al*., 2013, Schudt *et al*., 2015, Becker *et al*., 1996, Kolesnikova *et al*., 2004a, Kolesnikova *et al*., 2004b). According to these reports, nucleocapsid formation occurs in the perinuclear inclusion bodies, following which they are transported into the cytoplasm and redistributed prior to budding through the cell surface. The velocity of nucleocapsid movement inside cells ranges from 100 nm/s to 500 nm/s in MARV- or EBOV-infection (Schudt et al., 2013, Schudt et al., 2015). Application of specific cytoskeleton inhibitors revealed that the transport of MARV nucleocapsids was dependent on actin polymerization. Viral matrix protein VP40, which is a peripheral membrane protein and plays a pivotal role in filamentous virion formation, is essential for the recruitment of nucleocapsids to the cell periphery and for their incorporation into progeny virions (Noda *et al*., 2007, Bharat et al., 2012, Noda *et al*., 2002, Dolnik *et al*., 2010). The surface glycoprotein GP, which is an integral membrane protein and is responsible for cell entry, forms the filamentous virions together with VP40 (Fig. 1b) (Becker et al., 1996, Mittler *et al*., 2007, Beniac & Booth, 2017, Booth *et al*., 2013).

**Figure 1.**
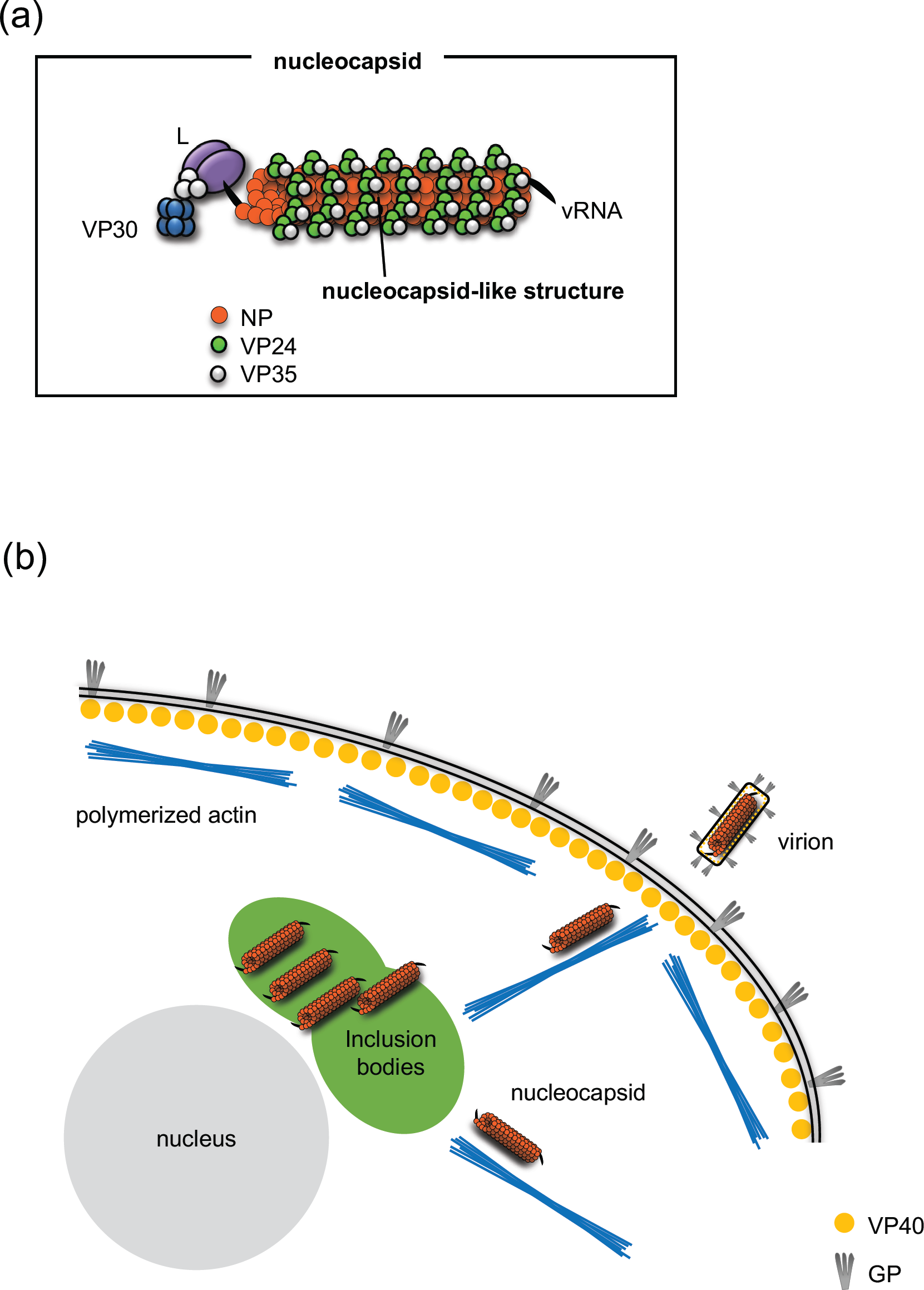
Orientation of MARV nucleocapsid and its transport pathway. (a) The viral genome is encapsidated by NP. Nucleocapsids additionally contain VP24, VP30, VP35, and L. Among them, NP, VP24, and V35 form the core structure of the nucleocapsid called a “nucleocapsid-like structure (NCLS)”. (b) Nucleocapsids are formed in the perinuclear inclusion bodies and are subsequently transported along polymerized actin filaments to the plasma membrane, where budding and release of virions takes place. The VP40 protein forms filamentous virions together with GP.

Live-cell imaging is a powerful tool for visualization of the spatio-temporal dynamics of living organisms. In addition to immunofluorescence microscopy,live-cell imaging microscopy has been utilized to visualize the localization of viral proteins, and interactions between viral and host proteins in various virus-infected cells (Nanbo *et al*., 2013, Hoenen *et al*., 2012, Lakdawala *et al*., 2014, Becker et al., 1996). However, because of its high pathogenicity, MARV must be handled under the highest biosafety conditions [biosafety level 4 (BSL-4)], which complicates and delays research using live-cell imaging (Falzarano *et al*., 2011). In this study, we developed a safe, live-cell imaging system, following a previously established method for EBOV (Takamatsu *et al*., 2018), to visualize MARV nucleocapsid-like structures (NCLSs) in cells expressing viral proteins, outside of BSL-4 laboratories. By using this live-cell imaging system, we were able to analyze interactions between NCLSs and the cellular cytoskeleton, as well as intracellular transport of NCLSs.

## Materials and methods

### Cell culture

Huh-7 (human hepatoma) cells were maintained at 37 °C and 5% CO_2_ in Dulbecco’s Modified Eagle Medium (DMEM, Life Technologies) supplemented with 10% (vol/vol) Fetal bovine serum (FBS, PAN Biotech), 5 mM L-glutamine (Q; Life Technologies), 50 U/mL penicillin, and 50 μg/mL streptomycin (PS; Life Technologies).

### Plasmids and transfection

Plasmids encoding the MARV structural proteins (NP, VP35, VP24, L, VP40 and GP): pCAGGS-NP, pCAGGS-VP35, pCAGGS-VP24, pCAGGS-L, pCAGGS-VP40, and pCAGGS-GP, and a MARV minigenome-expressing plasmid which encodes a *Renilla* luciferase were used (Wenigenrath *et al*., 2010, Hoenen *et al*., 2011). The plasmid pCAGGS-VP30-GFP, coding for the green fluorescent protein-VP30 fusion protein, was produced as previously described (Schudt et al., 2013). The transfection was performed in 50 μL Opti-MEM without phenol red (Life Technologies) using TranSIT (Mirus) according to the manufacturer’s instructions.

### Lice cell imaging microscopy

A total of 2×10^4^ Huh-7 cells were seeded onto a µ-Slide 4 well (Ibidi) and cultivated in DMEM/PS/Q with 10% FBS. Each well was transfected with the following plasmids, encoding all MARV structural proteins: (250 ng of pCAGGS-NP, 50 ng of pCAGGS-VP35, 125 ng of pCAGGS-VP30-GFP, 50 ng of pCAGGS-VP24, 500 ng of pCAGGS-L, 125 ng of pCAGGS-VP40 and 125 ng of pCAGGS-GP), together with a T7-driven, MARV minigenome-expressing plasmid, which encodes a *Renilla* luciferase, and a T7 polymerase-coding plasmid (pCAGGS-T7) (Wenigenrath et al., 2010, Hoenen et al., 2011). The inoculum was removed at 1 h post-transfection (p.t.), and 500 μL CO_2_-independent Leibovitz’s medium (Life Technologies) with PS/Q, non-essential amino acid solution, and 20% (vol/vol) FBS were added. Live-cell time-lapse experiments were recorded with a Nikon ECLIPSE TE2000-E using a 63× oil objective or a GE healthcare Delta Vision Elite using a 60× oil objective in biosafety level-2 laboratories.

### Treatment of cells with cytoskeleton-modulating drugs

Cells were treated with 15 μM nocodazole (Sigma), 0.3 μM cytochalasin D (Sigma), or 0.15% dimethyl sulfoxide (DMSO, Sigma), following previous publication (Schudt et al., 2013). The chemicals were added to the cell culture medium 3 hours prior to observation.

### Image processing and analysis

Acquired pictures and movie sequences were processed using the Fiji plugin “TrackMate” (Tinevez *et al*., 2017, Schindelin *et al*., 2012).

## Results and Discussion

### Establishment of a live-cell imaging system for MARV NCLSs transport

The Marburg virus VLP system, which models a complete, single infectious cycle, has been developed and used to analyze transcription and replication, as well as the budding processes (Wenigenrath et al., 2010, Hoenen et al., 2011). In this study, we attempted to visualize MARV NCLSs transport in Huh-7 cells, which is a suitable method to determine the sequential intracytoplasmic movement of MARV nucleocapsids (Schudt et al., 2013), by using this VLP system following the procedure established in EBOV (Takamatsu et al., 2018). Transfection of the VLP components with the following modifications: replacing VP30 with VP30-GFP, was performed as described in Figure 2a. We employed this VLP-based system in all subsequent experiments in this study.

**Figure 2.**
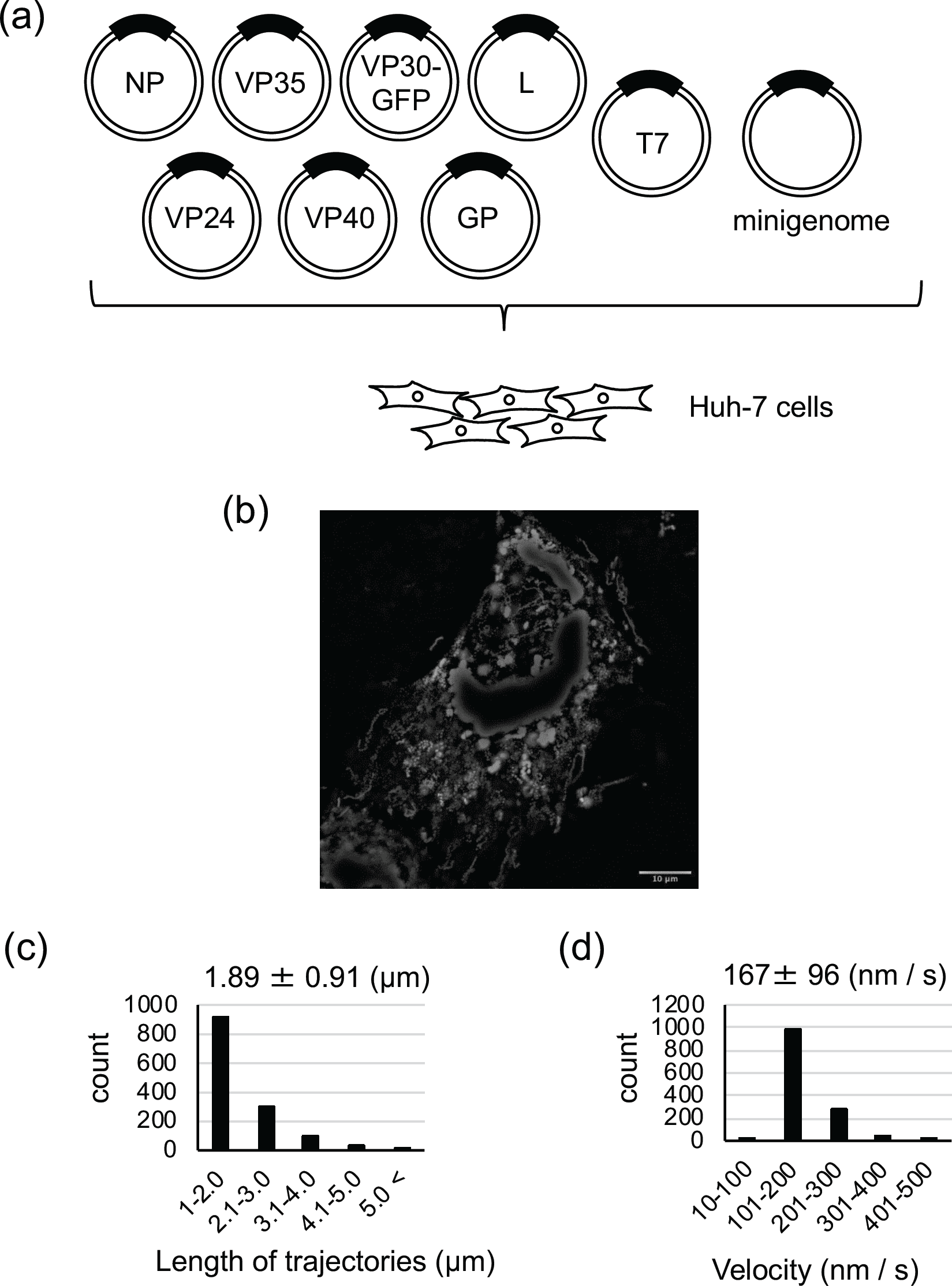
Live-cell imaging system of MARV nucleocapsid-like structures. (a) The experimental setting for detection of NCLS transport. Huh-7 cells were transfected with plasmids encoding NP, L, VP35, VP24, VP40, GP, Marburg virus specific minigenome, T7 polymerase, and VP30-GFP. (b) Plasmid-transfected Huh-7 cells were observed at 18 h p.t. The image shows the maximum-intensity projection of time-lapse images of cells, recorded for 90 seconds; images were captured every 2 s. (c, d) Over 100,000 moving signals were captured and over 1,000 selected signals were analyzed using the Fiji plugin “TrackMate”. (c) The length of the NCLS trajectories was evaluated. The y-axis represents the number of signals in each range (x-axis). The numbers indicate mean ± SD (μm). (d) The velocity of NCLSs transport was evaluated. The y-axis represents the number of signals in each range (x-axis). The numbers indicate mean ± SD (nm/s).

At 18 h p.t., we detected a large number of signals with various shapes (Supplementary Movie 1). Over 1,000 signals in the acquired movie sequences were analyzed. The representing sequence was expressed as the maximum intensity projection, in which the image projected maximum intensity of each time-lapse were plotted. (Fig. 2b). Among the signals, we focused on the signals showing long-distance and directional transport, which represent NCLSs trajectories (Takamatsu et al., 2018). We detected trajectories with lengths ranging from 100 nm to 20 µm, with a mean length of 1.89 ± 0.91 μm (Fig. 2c). The direction of movement of each NCLS also varied, though the cause of this variation remains unclear. The speed of MARV NCLSs transport ranged from 10 nm/s to 500 nm/s, with a mean velocity of 167 ± 96 nm/s (Fig. 2d), which is comparable to the velocity of nucleocapsids transport in MARV-infected cells analyzed in BSL-4 laboratories (Schudt et al., 2013). In summary, the movement characteristics of NCLSs in our system is similar to those of nucleocapsids in MARV-infected cells.

### Actin polymerization is required for MARV NCLSs transport

In MARV-infected cells, the microtubule depolymerizing drug nocodazole does not alter the movement of nucleocapsids, whereas the actin depolymerizing drug cytochalasin D arrests it (Schudt et al., 2013). To confirm the relevance of the live-cell imaging system we developed in this study, we analyzed NCLSs movement after treatment with cytoskeletal modulation drugs. Huh-7 cells were transfected with plasmids as described in Figure 2a. The culture medium was replaced at 15 h p.t. with Leibovitz’s medium containing either 0.15 % DMSO (control), 0.15 M nocodazole, or 0.3 µM cytochalasin D (Fig. 3a-d, Supplementary Movie 2-4) (Schudt et al., 2013). After incubation the cells with cytoskeletal modulating drugs for 3h, time-lapse images were acquired. Nocodazole treatment did not alter the trajectory length of NCLSs transport in comparison to the control, whereas cytochalasin D treatment induced immediate cessation of long-distance transport (Fig. 3e-g). The mean velocity of NCLSs transport in the control or nocodazole-treated cells was 188 ± 98 nm/s and 219 ± 99 nm/s, respectively (Fig. 3h). On the other hand, only a few non-specific NCLSs movements were detectable in cytochalasin D-treated cells, with a mean velocity of 2.73 ± 23 nm/s (Fig. 3i-j). These results confirmed that NCLSs transport is dependent on actin polymerization, as well as the availability of our assay to test candidate drugs which target nucleocapsid intracellular transport.

**Figure 3.**
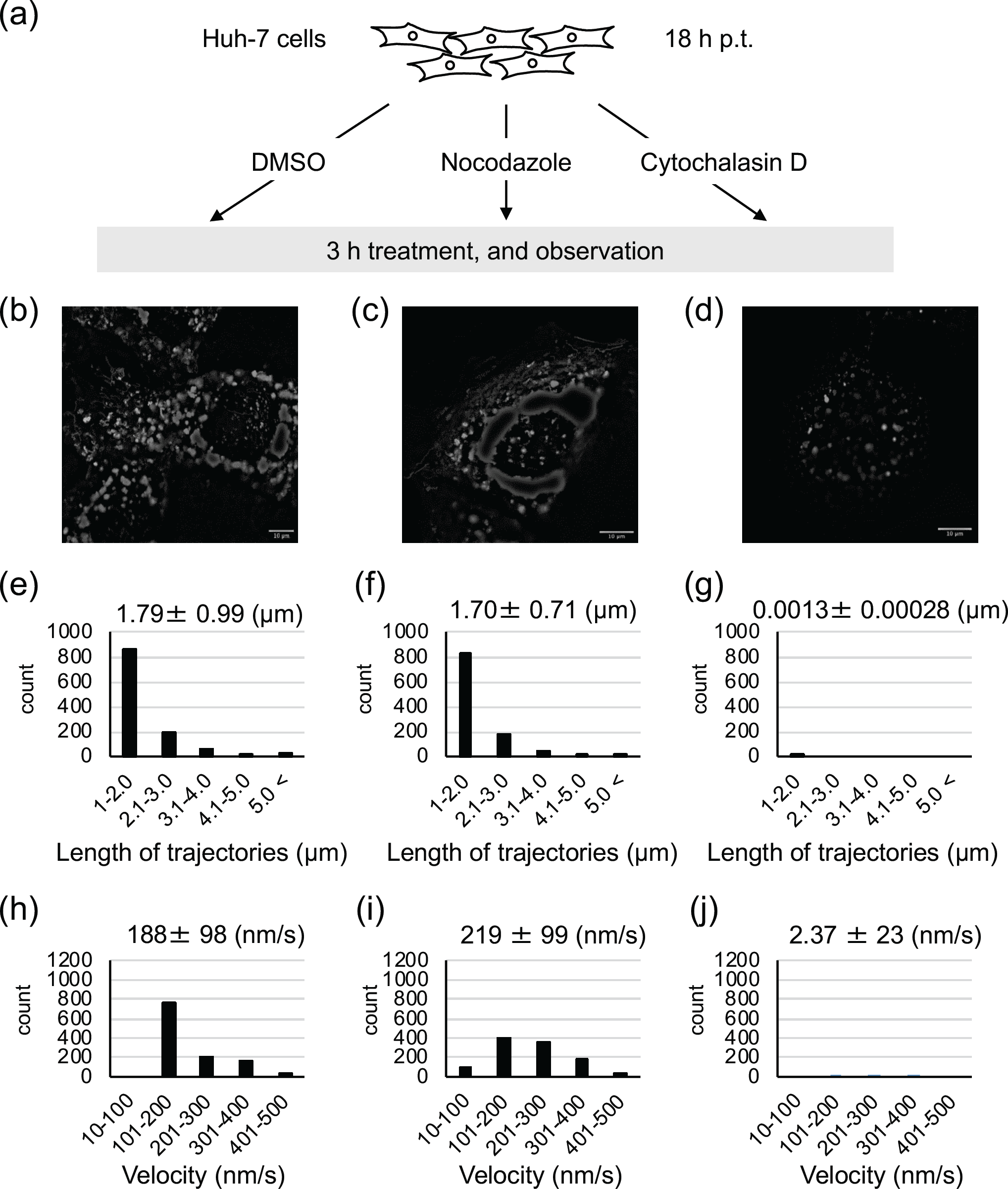
Effect of cytoskeleton-modulating drugs on MARV NCLSs transport. (a) Experimental setting to observe the effects of cytoskeleton modulating drugs on MARV NCLSs transport. Huh-7 cells were transfected with plasmids as described in Fig. 2(a), and treated at 15 h p.t. with either 0.15 % DMSO (control), 0.15 μM nocodazole, or 0.3 μM cytochalasin D. After 3 hours of treatment, observation of the cells began (18 h p.t.). (b-d) Time-lapse images were acquired for each of the drug treated cells (b: DMSO, c: Nocodazole, d: Cytochalasin D). The pictures show the maximum-intensity projection of time-lapse images of cells, recorded for 90 seconds; images were captured every 2 s. (e-i) Over 100,000 moving signals were captured and over 1,000 selected signals were analyzed using Fiji plugin “TrackMate”. (e-g) The length of the NCLS trajectories was evaluated in each of the drug treated cells (e: DMSO, f: Nocodazole, g: Cytochalasin D). The y-axis represents the number of signals in each range (x-axis). The numbers indicate mean ± SD (μm). (h-i) The velocity of NCLSs transport was evaluated in each of the drug treated cells (h: DMSO, i: Nocodazole, j: Cytochalasin D). The y-axis represents the number of signals in each range (x-axis). The numbers indicate mean ± SD (nm/s).

### Concluding remarks

In the present study, we developed a live-cell imaging system for cells expressing viral proteins, which can be safely used in BSL-2 laboratories. Furthermore, we demonstrated the relevance of our system as a substitute for the analysis of nucleocapsids transport in MARV-infected cells. Currently, cellular factors involved in the nucleocapsid transport have not been fully understood. The combined approach of gene silencing and inhibitor screening using the system we developed can be utilized to identify the key host factors for the intracellular transport of MARV nucleocapsids. Moreover, it is noteworthy that the technical approach developed here might be applicable to study the nucleocapsid transport of other mononegaviruses, as well as to characterize antivirals inhibiting nucleocapsid transport.

## Acknowledgments

The authors are grateful to Olga Dolnik and Gordian Schudt (Philipps University Marburg, Germany) for fruitful discussion.

